# 5WBF: A low-cost and straightforward whole blood filtration method suitable for whole-genome sequencing of *Plasmodium falciparum* clinical isolates

**DOI:** 10.1101/2021.08.30.457783

**Authors:** Romain Coppée, Atikatou Mama, Véronique Sarrasin, Claire Kamaliddin, Lucie Adoux, Lawrence Palazzo, Nicaise Tuikue Ndam, Franck Letourneur, Frédéric Ariey, Sandrine Houzé, Jérôme Clain

## Abstract

**Background:** Whole-genome sequencing (WGS) is becoming increasingly helpful to assist malaria control programs. A major drawback of this approach is the large amount of human DNA compared to parasite DNA extracted from unprocessed whole blood. As red blood cells (RBCs) have a diameter of about 7-8 μm and exhibit some deformability, we hypothesized that cheap and commercially available 5 μm filters might retain leukocytes but much less of *Plasmodium falciparum*-infected RBCs. This study aimed to test the hypothesis that such a filtration method, named 5WBF (for <u>5</u> μm <u>W</u>hole <u>B</u>lood <u>F</u>iltration), may provide highly enriched parasite material suitable for *P. falciparum* WGS.

**Methods:** Whole blood was collected from five patients experiencing a *P. falciparum* malaria episode (ring-stage parasitemia range: 0.04-5.5%) and from mock samples obtained by mixing synchronized, ring-stage cultured *P. falciparum* 3D7 parasites with uninfected human whole blood (final parasitemia range: 0.02-1.1%). These whole blood samples (50 to 400 μL) were diluted in RPMI 1640 medium or PBS 1X buffer and filtered with syringes connected to a 5 μm commercial filter. DNA was extracted from filtered and unfiltered counterpart blood samples using a commercial kit. The 5WBF method was evaluated on the ratios of parasite:human DNA assessed by qPCR and by sequencing depth and percentages of coverage from WGS data (Illumina NextSeq 500). As a comparison, we also applied to the same unprocessed whole blood samples the selective whole-genome amplification (sWGA) method which does not rely on blood filtration.

**Results:** After applying 5WBF, qPCR indicated an average of 2-fold loss in the amount of parasite template DNA (Pf ARN*18S* gene) and from 4,096- to 65,536-fold loss of human template DNA (human *β actin* gene). WGS analyses revealed that > 95% of the nuclear genome and the entire whole organellar genomes were covered at ≥ 10× depth for all samples tested. In sWGA counterparts, none of the organellar genomes were covered, and from 47.7 to 82.1% of the nuclear genome was covered at ≥ 10× depth depending on parasitemia. Sequence reads were homogeneously distributed across gene sequences for 5WBF-treated samples (n = 5,460 genes; mean coverage: 91×; median coverage: 93×; 5^th^ percentile: 70×; 95^th^ percentile: 103×), allowing the identification of gene copy number variations such as for *gch1*. This later analysis was not possible for sWGA-treated samples, as we observed a much more heterogeneous distribution of reads among gene sequences (mean coverage: 80×; median coverage: 51×; 5^th^ percentile: 7×; 95^th^ percentile: 245×).

**Conclusions:** The novel 5WBF leucodepletion method is simple to implement and based on commercially available, standardized, 5 μm filters which cost from 1.0 to 1.7€ per unit, depending on suppliers. 5WBF permits extensive genome-wide analysis of *P. falciparum* DNA from minute amounts of whole blood even with parasitemias as low as 0.02%.

## INTRODUCTION

Whole-genome sequencing (WGS) has revolutionized genome-wide analyses [1]. In the context of *Plasmodium falciparum* surveillance, WGS is helpful for example to analyze the structure of parasite populations [2,3] and to identify and track gene mutations conferring resistance to antimalarial drugs [4,5]. A major drawback of WGS is the large amount of human DNA compared to parasite DNA when extracted from unprocessed whole blood. Several protocols have been developed to enrich parasite DNA before WGS, either by filtering out leukocytes before DNA extraction or by selectively amplifying the parasite genome (sWGA). Current filtration procedures based on leucodepletion are however limited because they require large volumes of venous blood [6] or use either home-made cellulose-packed columns [7] or costly commercial filters [8]. Regarding sWGA-based methods [9–11], studies reported a large proportion of unmapped reads to the *P. falciparum* genome [12], the absence of coverage of the organellar genomes [9], and a wide heterogeneity in read distribution across the nuclear genome [9]. Altogether, there is a need to improve clinical sample preparation to increase data quality and exhaustiveness for *P. falciparum* genomic studies while limiting the cost of data production.

Mature red blood cells (RBCs) have a resting long diameter of about ~8 μm and exhibit some deformability [13]. Using a microfluidic device to examine the traversal of a RBC, the diameters of the smallest equivalent cylindrical tube, through which uninfected and parasitized RBCs could pass, were similar (2.78 and 2.79 respectively) [13]. Human leukocytes are larger cells than RBCs and have diameters ranging from 9 to 21 μm depending on cell types. Hence, we hypothesized that commercially available filters with a pore size of 5 μm might retain DNA-carrying human leukocytes but not *P. falciparum*-infected RBCs. Such a filtration could provide samples highly enriched in parasites, suitable for downstream WGS workflow. On this basis, we developed 5WBF (<u>5</u> μm <u>W</u>hole <u>B</u>lood <u>F</u>iltration), a low-cost and simple blood filtration procedure using a commercial, standardized 5 μm filter (Minisart NML® syringe, Sartorius AG, Germany). To develop 5WBF, 400 μL of whole blood at variable parasitemias (from 0.022 to 1.1%) was first tested from mock samples made by mixing synchronized, ring-stage cultured *P. falciparum* parasitized erythrocytes (3D7) with uninfected human whole blood. Then 5WBF was validated using 50 and 200 μL of whole blood (ring-stage parasitemia range: 0.04-5.5%) from patients experiencing a *P. falciparum* malaria episode. DNA extracts obtained after 5WBF were then evaluated using the parasite:human DNA ratio assessed by qPCR and the performance of sequencing depth and percentages of coverage obtained through WGS data compared with sWGA.

## MATERIALS AND METHODS

### *P. falciparum* culture and infected whole blood reconstitution

Mock whole blood samples were obtained by mixing a synchronized ring-stage *P. falciparum* culture (O-negative blood, 3D7 parasite strain) with uninfected human whole blood (final parasitemia range: 0.022-1.1%). 3D7 parasites were cultured at 37°C under specific atmospheric conditions (10% oxygen, 5% carbon dioxide and 85% nitrogen) in 10% human serum containing RPMI 1640 medium. One volume of pelleted culture at 10% parasitemia was diluted in ten volumes of non-infected human whole blood. The mock sample parasitemia was then estimated to be 1.1% by Diff Quick™-stained thin blood film. The sample was then diluted 1:5 followed by another 1:10 dilution in the negative human whole blood as three independent replicates. The parasitemias were then estimated by Diff Quick™-stained thin blood film to be 0.23% and 0.022% for the two diluted samples, respectively. These reconstituted, infected whole blood samples were hereafter called mock samples.

### Infected whole blood from *P. falciparum* malaria patients

Five fresh blood samples (collected on EDTA) from imported *P. falciparum* malaria cases diagnosed and treated at Bichat-Claude Bernard Hospital (French Malaria Reference Center, Paris, France) were arbitrarily selected. Diff Quick™-stained blood film examination indicated monospecific *P. falciparum* infections with ring-stage parasitemias ranging from 0.04 to 5.5%. Additional clinical information is provided for each patient in **Additional file 1: Table S1**.

### 5WBF procedure

Prior to filtration, whole blood was diluted in either PBS 1X buffer or RPMI 1640 medium (**Figure 1** and **Table 1**). For clinical samples, 50 μL and 200 μL of whole blood were diluted in 30 volumes of RPMI 1640 medium, as using small sample volumes would result in large sample loss. For mock samples, 400 μL of whole blood were diluted in ten volumes of PBS 1X buffer (**Table 1**). Each diluted blood sample was then filtered using a 5 μm surfactant-free cellulose acetate syringe filter (Minisart NML® syringe filter, Sartorius reference number: 17594K) connected to a 10 mL syringe (**Figure 1**). The sample was filtered by a very gentle push with the syringe plunger such that the filtrate flew drop by drop. Importantly, the plunger was pushed to the bottom of the syringe. Note that even though the filtrate passes by gravity only, using the plunger is seemingly mandatory to recover a maximum of infected RBCs. For mock samples only, the filter membrane was rinsed with another 2 mL of PBS 1X buffer (**Table 1**). The filtrate was then centrifuged at 2,500 *g* for 5 minutes at room temperature and the supernatant was discarded (**Figure 1**). One pellet volume of RPMI 1640 (clinical samples) or PBS 1X (mock samples) was added to the pelleted RBCs which were transferred to 2 mL tube and stored at 4°C until DNA extraction within the next 24 hours (**Table 1**). The filtration step itself is very quick (1 to 3 minutes), and the whole procedure takes about 20 minutes.

**Table 1.** Details of samples subjected to WGS.

| Origin | Sample | Enrichment method | % par.<sup>a</sup> | Blood volume (μL) | Blood:buffer dilution | Washing filter membrane<sup>b</sup> | Volume of DNA elution buffer (μL) |
| --- | --- | --- | --- | --- | --- | --- | --- |
| Mock (3D7) | M1 | none | 1.10 | 400 | 1:10, PBS 1X | n.a. | 200 |
|  | M1<sub>WGA</sub> | sWGA | 1.10 | 400 | 1:10, PBS 1X | n.a. | 200 |
|  | M1-1<sub>5F</sub> | 5WBF | 1.10 | 400 | 1:10, PBS 1X | 2 mL PBS 1X | 200 |
|  | M1-2<sub>5F</sub> | 5WBF | 1.10 | 400 | 1:10, PBS 1X | 2 mL PBS 1X | 200 |
|  | M1-3<sub>5F</sub> | 5WBF | 1.10 | 400 | 1:10, PBS 1X | 2 mL PBS 1X | 200 |
|  | M2<sub>WGA</sub> | sWGA | 0.23 | 400 | 1:10, PBS 1X | n.a. | 200 |
|  | M2-1<sub>5F</sub> | 5WBF | 0.23 | 400 | 1:10, PBS 1X | 2 mL PBS 1X | 200 |
|  | M2-2<sub>5F</sub> | 5WBF | 0.23 | 400 | 1:10, PBS 1X | 2 mL PBS 1X | 200 |
|  | M2-3<sub>5F</sub> | 5WBF | 0.23 | 400 | 1:10, PBS 1X | 2 mL PBS 1X | 200 |
|  | M3<sub>WGA</sub> | sWGA | 0.022 | 400 | 1:10, PBS 1X | n.a. | 200 |
|  | M3-1<sub>5F</sub> | 5WBF | 0.022 | 400 | 1:10, PBS 1X | 2 mL PBS 1X | 200 |
|  | M3-2<sub>5F</sub> | 5WBF | 0.022 | 400 | 1:10, PBS 1X | 2 mL PBS 1X | 200 |
|  | M3-3<sub>5F</sub> | 5WBF | 0.022 | 400 | 1:10, PBS 1X | 2 mL PBS 1X | 200 |
| Patients | P1<sub>5F-50</sub> | 5WBF | 0.04 | 50 | 1:30, RPMI 1640 | n.d. | 200 |
|  | P1<sub>5F-200</sub> | 5WBF | 0.04 | 200 | 1:30, RPMI 1640 | n.d. | 200 |
|  | P2<sub>5F-50</sub> | 5WBF | 0.08 | 50 | 1:30, RPMI 1640 | n.d. | 200 |
|  | P2<sub>5F-200</sub> | 5WBF | 0.08 | 200 | 1:30, RPMI 1640 | n.d. | 200 |
|  | P3<sub>5F-50</sub> | 5WBF | 0.25 | 50 | 1:30, RPMI 1640 | n.d. | 200 |
|  | P3<sub>5F-200</sub> | 5WBF | 0.25 | 200 | 1:30, RPMI 1640 | n.d. | 200 |
|  | P4<sub>5F-50</sub> | 5WBF | 0.40 | 50 | 1:30, RPMI 1640 | n.d. | 200 |
|  | P4<sub>5F-200</sub> | 5WBF | 0.40 | 200 | 1:30, RPMI 1640 | n.d. | 200 |
|  | P5<sub>5F-50</sub> | 5WBF | 5.50 | 50 | 1:30, RPMI 1640 | n.d. | 200 |
|  | P5<sub>5F-200</sub> | 5WBF | 5.50 | 200 | 1:30, RPMI 1640 | n.d. | 200 |

Note – <sup>a</sup> % par., parasitemia in percentage; <sup>b</sup> n.a., not applicable; n.d., not done

**Figure 1.**
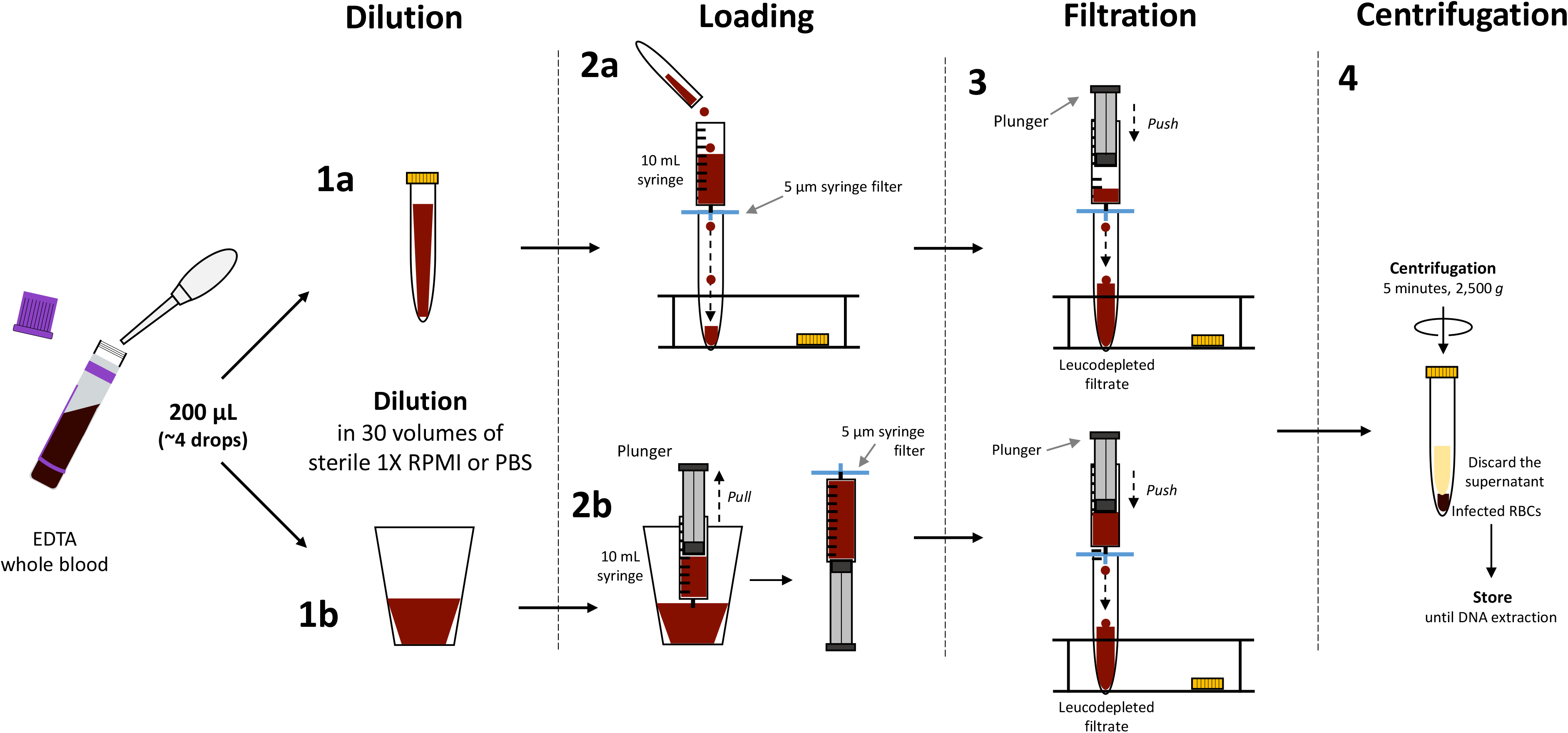
Main steps of 5WBF. (**1**) 50 to 400 μL of whole blood were diluted in RPMI 1640 medium or PBS 1X buffer in either (a) a 15 mL tube or (b) a larger flask. The cartoon shows 200 μL of whole blood diluted in RPMI 1640 as an example. (**2**) The diluted sample was loaded onto a 10 mL syringe either after (a) or before (b) the 5 μm filter was connected to the syringe. (**3**) The blood was filtered by very gentle pressure with the syringe plunger. The pressure was applied until the plunger reached the bottom of the syringe to recover the maximum of infected RBCs. The filtration step itself is rapid and takes about 1 to 3 minutes. (**4**) The filtrate was centrifuged at 2,500 *g* for 5 minutes and the supernatant was discarded. The pellet was suspended with ~ one volume of RPMI 1640 or PBS 1X, transferred into a 1.5 mL tube, and stored until DNA extraction. Important notes: *i*) from our own experiments, we estimated that the filter dead volume was about 200 μL (reported as 100-150 μL by the manufacturer); *ii*) even after the 2 mL optional wash, the filter had a red color indicating some retained RBCs or hemolysis during filtration; RBCs loss seems low although we have not precisely quantified it; *iii*) the harder the push with the syringe plunger, the more hemolysis occurs; and *iv)* the WGS data presented in this study were obtained using the (2b) version of the protocol; similar high-quality WGS data were obtained when using the (2a) version (data not shown).

As a negative control for filtration, whole blood (400 μL) for one mock sample (parasitemia of 1.1%) was subjected to the same pipeline similarly to other mock samples, except that we omitted to connect a filter at the bottom of the syringe (**Table 1**). This control mock sample is latter called M1.

### DNA extraction

DNA extraction was carried out on unfiltered and 5WBF-treated samples using the MagPurix® Blood DNA Extraction Kit 200 (Biosynex, France), then eluted using the elution buffer according to the manufacturer’s recommendations (**Table 1**). DNA was quantified using the Qubit® dsDNA high sensitivity kit (Thermo Fisher Scientific) according to the manufacturer’s recommendations.

### sWGA procedure

The sWGA method was performed on genomic DNA from unfiltered mock samples according to published protocols [9]. The sWGA reaction was performed in 0.2 mL PCR-tubes, containing 10 ng of template genomic DNA, 1× BSA (New England Biolabs), 1 mM dNTPs (New England Biolabs), 2.5 μM of each amplification primer (**Additional file 1: Table S2**), 1× Phi29 reaction buffer (New England Biolabs), 30 units of Phi29 polymerase (New England Biolabs), and molecular biology grade water to reach a final reaction volume of 50 μL. The reaction was carried out on a thermocycler with the following step-down program: 5 min at 35°C, 10 min at 34°C, 15 min at 33°C, 20 min at 32°C, 30 min at 31°C, 16 h at 30°C, then heating for 15 min at 65°C to inactivate the Phi29 polymerase before cooling to 4°C. Amplified products were quantified using the Qubit® dsDNA high sensitivity kit (Thermo Fisher Scientific) to determine whether there was at least 500 ng of product for sequencing. Amplified products were cleaned using Agencourt Ampure XP beads (Beckman Coulter) as follows: 1.8 volumes of beads were added to 1 volume of amplified products, briefly mixed, and then incubated for 5 min at room temperature. A magnetic rack was used to capture the DNA binding beads. The DNA binding beads were then washed twice using 200 μL of 80% ethanol and eluted with 60 μL of EB buffer.

### Quantitative PCR

The copy number of specific *P. falciparum* and human genes within the genomic DNA from clinical samples was estimated by qPCR with *Plasmodium* Typage kit (Bio-Evolution, France). Briefly, as recommended by the manufacturer, 5 μL of DNA extract was mixed with 15 μL of Master Mix containing specific primers and probes targeting *P. vivax* and *P. falciparum* ARN*18S* and human *β actin* genes. The reaction was carried out on a thermocycler (ViiA 7, Applied Biosystems) with the following program: 30 sec at 95°C; 40 cycles: 15 sec at 95°C followed by 45 sec at 60°C; then 1 sec at 37°C. Positive and negative controls were included in each run.

### Whole-genome sequencing

Genomic DNA libraries were constructed for high throughput sequencing using the KAPA hyper Prep Library Preparation Kit (Kapa Biosystems, Woburn, MA). Mechanical DNA shearing was performed with the Covaris S220 through microTube-50 AFA Fiber Screw-Cap (Covaris®) using a setting of 30% duty factor, 100W peak incidence power, and 1000 cycles per burst for 150 seconds. Then, DNA libraries were checked for quality and quantity using Qubit® for concentration and BioAnalyser 2100 Agilent for fragment size. Libraries were sequenced at 150 bp paired-end using an Illumina NextSeq 500 instrument at the GENOM’IC platform from Institut Cochin (Paris, France).

### Sequencing output analysis

Sequence data obtained from each sample was subjected to standard Illumina QC procedures. Each sample was analyzed independently by mapping sequence reads to the *P. falciparum* 3D7 reference genome v.39 using the BWA software package [14]. Samtools (http://samtools.sourceforge.net/) was used to generate coverage statistics and depth estimates from the BWA mapping output. Qualimap v2.2.1 was also used to perform an analysis based on specific features derived from the alignment, including coverage, GC content and mapping quality [15]. A home-made python script was developed to calculate the percentage of each *P. falciparum* gene covered at ≥ 10× depth and the per-gene mean coverage depth (https://github.com/Rcoppee/Scan_gene_coverage). The script required a reference genome file in fasta format, an annotation gff file indicating the location of each exon, intron and corresponding genes, and a per-base coverage depth file generated with Bedtools genomecov function using default parameters [16]. This per-base coverage depth file was also used to plot the average read depth in 1 kb windows across the 14 *P. falciparum* chromosomes using the Circos software [17]. Finally, per-gene copy number was assessed using PlasmoCNVScan, a custom read depth strategy specifically made for *Plasmodium* species [18].

### Ethical considerations

Samples received at the French Malaria Reference Center (Paris, France) were registered and declared for research purposes as a biobank for both the Assistance Publique des Hôpitaux de Paris and Santé Publique France. The uninfected blood sample was obtained from a patient having a negative malaria test. No institutional review board approval was required according to French legislation (article L. 1111-7 du Code de la Santé Publique, article L. 1211-2 du Code de Santé Publique, articles 39 et suivants de la loi 78-17 du 6 janvier 1978 modifiée en 2004 relative à l’informatique, aux fichiers, et aux libertés).

## RESULTS

### Application of 5WBF to mock samples

5WBF was first applied to 400 μL of mock blood samples (3D7 culture diluted in uninfected whole blood) with parasitemias of 1.1, 0.23 and 0.022% (each in triplicate). For the unfiltered control mock sample (sample M1; 1.1% parasitemia), 21.2% of the reads mapped to the *P. falciparum* genome with a mean coverage of 5.6× depth (**Figure 2A** and **Table 2**). For the 5WBF-treated mock samples (n = 9), an average of 89.6% (standard deviation: 11.4) of the reads mapped to the *P. falciparum* genome. The proportion of *P. falciparum*-mapped reads decreased with parasitemia (**Figure 2A** and **Table 2**). Regardless, *P. falciparum* genome coverage at ≥ 10× depth was at least 98.6% whatever the parasitemia (**Figure 2B**). Reads covered both the parasite’s mitochondrial and apicoplast genomes, for which mean coverages were systematically higher than 1,000× and 59.6× depths respectively. For comparison, one unfiltered mock sample of each parasitemia was processed by the sWGA procedure. Higher proportions of reads mapped to the *P. falciparum* genome when produced from 5WBF-treated compared to sWGA-treated samples (**Figure 2A** and **Table 2**). The difference was modest at 1.1% parasitemia but it increased as parasitemias dropped to 0.23 and 0.022%. Also, reads from sWGA poorly covered the parasite mitochondrial and apicoplast genomes, systematically below a mean coverage of 10× depth (**Table 2**). In summary almost all the bases of the different genomes of *P. falciparum* were analyzable at ≥ 10× depth using 400 μL of 5WBF-treated whole blood.

**Table 2.** WGS statistics of *P. falciparum* DNA extracted from mock samples subjected to either no treatment or sWGA or 5WBF.

| Sample name | % par.<sup>a</sup> | Enrichment method | Total reads | *P. falciparum*-mapped reads (%) | Mean genome coverage (X) | Mean coverage of mt genome (X)<sup>b</sup> | Mean coverage of api genome (X)<sup>c</sup> |
| --- | --- | --- | --- | --- | --- | --- | --- |
| M1 | 1.1 | None | 14,777,245 | 3,135,321 (21.22) | 5.6 | 45.39 | 11.76 |
| M1<sub>WGA</sub> | 1.1 | sWGA | 16,002,711 | 15,347,302 (95.9) | 74.4 | 5.20 | 8.44 |
| M1-1<sub>5F</sub> | 1.1 | 5WBF | 19,932,109 | 19,558,272 (98.12) | 104.8 | 2102.17 | 216.15 |
| M1-2<sub>5F</sub> | 1.1 | 5WBF | 16,923,729 | 16,591,158 (98.03) | 89.59 | 1701.08 | 183.29 |
| M1-3<sub>5F</sub> | 1.1 | 5WBF | 17,230,853 | 17,018,367 (98.77) | 90.04 | 1335.70 | 150.66 |
| M2<sub>WGA</sub> | 0.23 | sWGA | 16,266,710 | 14,112,713 (86.76) | 69.16 | 1.59 | 3.57 |
| M2-1<sub>5F</sub> | 0.23 | 5WBF | 14,905,078 | 14,350,236 (96.28) | 79.71 | 1468.76 | 120.40 |
| M2-2<sub>5F</sub> | 0.23 | 5WBF | 19,310,196 | 18,169,487 (94.09) | 99.82 | 1782.77 | 161.75 |
| M2-3<sub>5F</sub> | 0.23 | 5WBF | 17,672,315 | 16,766,350 (94.87) | 91.15 | 1933.32 | 129.47 |
| M3<sub>WGA</sub> | 0.022 | sWGA | 9,840,798 | 4,915,197 (49.95) | 18.63 | 0.31 | 2.34 |
| M3-1<sub>5F</sub> | 0.022 | 5WBF | 17,283,136 | 14,467,900 (83.71) | 76.72 | 1209.77 | 115.57 |
| M3-2<sub>5F</sub> | 0.022 | 5WBF | 17,759,091 | 11,975,821 (67.43) | 62.32 | 1528.46 | 98.56 |
| M3-3<sub>5F</sub> | 0.022 | 5WBF | 13,052,530 | 9,839,100 (75.38) | 52.17 | 1026.41 | 59.63 |

Note – Reads were mapped to the *P. falciparum* 3D7 reference genome v.39. <sup>a</sup>% par., parasitemia in percentage; <sup>b</sup> mt, mitochondrial; <sup>c</sup> api, apicoplast.

**Figure 2.**
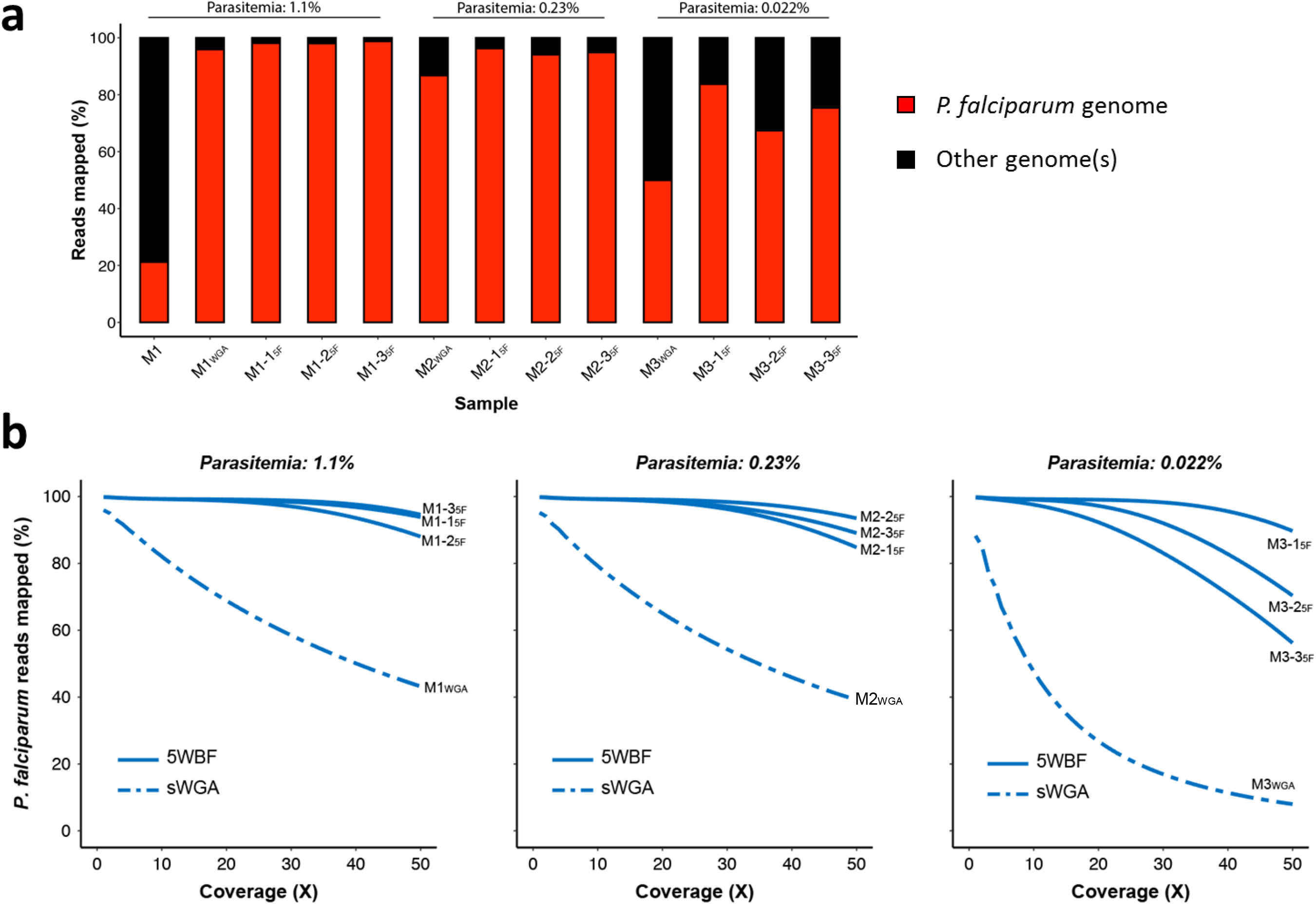
Proportions of mapped reads and *P. falciparum* genome coverage from mock whole blood samples. (**a**) Proportion of reads mapping to the *P. falciparum* nuclear and organellar genomes. Red and black colors indicate the proportion of reads mapping and not mapping to the *P. falciparum* genomes respectively. (**b**) Genome fraction coverage from 1× to 50× depth. Data from sWGA- and 5WBF-treated samples are indicated in dashed and solid lines respectively. Three independent 5WBF blood filtration replicates were made.

We then explored two gene-level metrics: the percentage of each *P. falciparum* gene covered at ≥ 10× depth and corresponding mean coverage. For these analyses, we used two samples presenting a similar number of reads mapping to the *P. falciparum* genome (M2_WGA_ and M2-1_5F_; parasitemia: 0.23%; ~14 million reads; **Table 2**).

First, for the 5WBF-treated sample, 99.0% (5,404/5,460) of nuclear genes and all organellar genes were fully covered at ≥ 10× depth, including important drug resistance genes such as *k13*, *mdr1*, *crt*, *dhfr* and *dhps* (**Figure 3A**). The few uncovered genes were mostly *rifin* and *var*. Using sWGA, 68.5% (3,741/5,460) of *P. falciparum* genes were fully covered at ≥ 10× depth. None of the mitochondrial and apicoplast genes nor the drug resistance gene *mdr1* were covered at ≥ 10× depth (**Figure 3B**).

**Figure 3.**
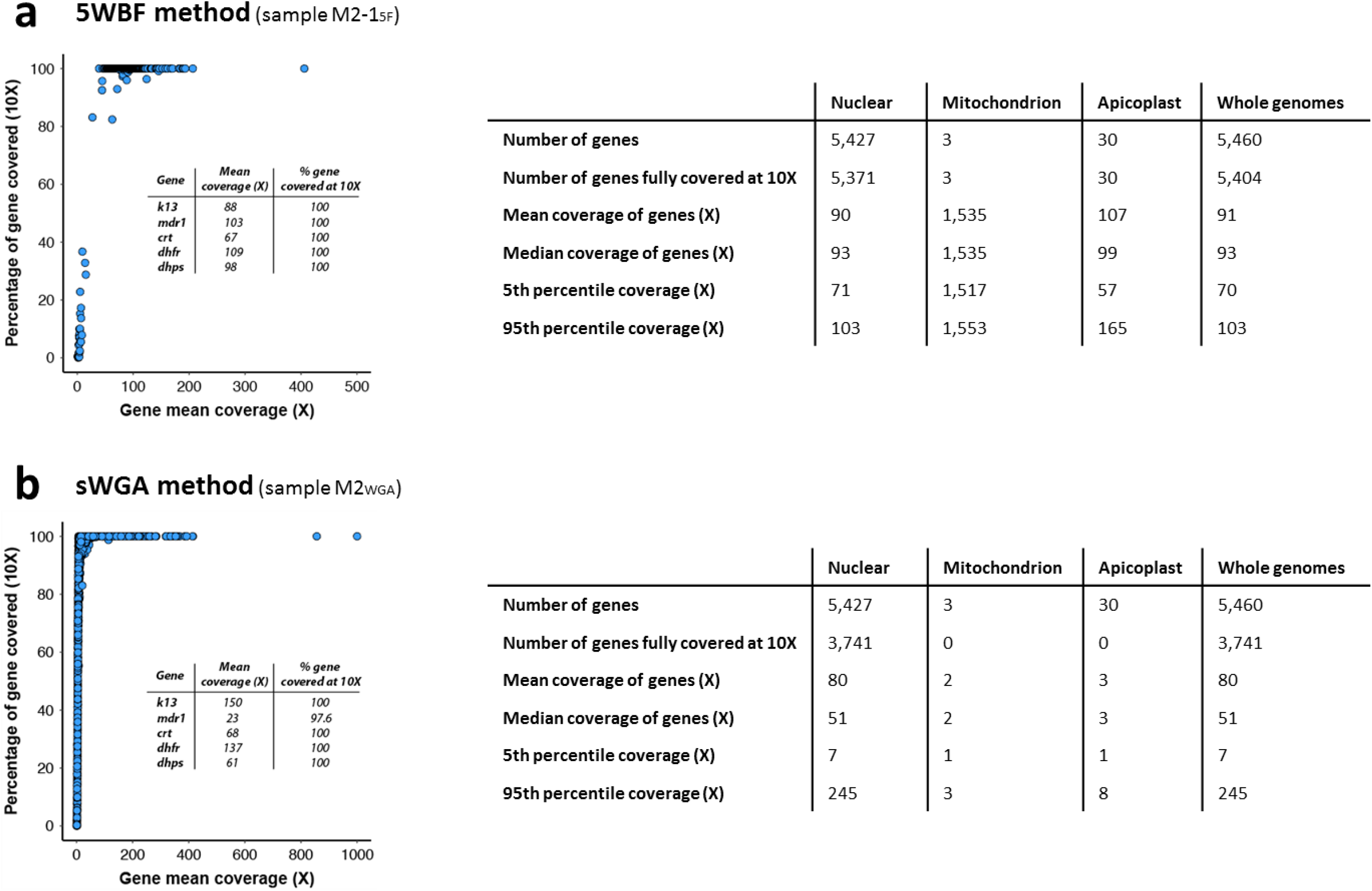
Comparison of gene coverage depth between M2_WGA_ and M2-1_5F_ mock samples. (**a**) Coverage depth of all genes for M2-1_5F_. Gene distribution based on the percentage of gene covered at ≥ 10× depth. Each blue point corresponds to a gene. Mitochondrial genes were discarded for ease of representation. The *insert* table indicates the mean coverage and the percentage of genes covered at ≥ 10× for five drug resistance genes. Descriptive statistics on the right of the figure included the number of genes, the number of genes fully covered at ≥ 10×, the mean and median coverage of all genes, and the 5^th^ and 95^th^ percentiles of coverage depth. Genes were partitioned as of either nuclear, mitochondrial, or apicoplast origins. (**b**) Coverage depth of all genes for M2_WGA_. Description of the plot and the tables are the same as in (a).

Second, the per-gene coverage varied little extensively with 5WBF (mean coverage: 90×; median: 93×; 5^th^ percentile: 71×; 95^th^ percentile: 103×) compared with the sWGA counterpart (mean coverage: 80×; median: 51×; 5^th^ percentile: 7×; 95^th^ percentile: 245×) (**Figure 3A** and **3B**). The coverage depth was also measured at each base across the 14 *P. falciparum* chromosomes. Reads mapped homogeneously across the genome with 5WBF, while they mapped much more heterogeneously in sWGA (**Figure 4**). Consequently, 5WBF is likely compatible with analyses based on read distribution such as identifying per-gene copy number.

**Figure 4.**
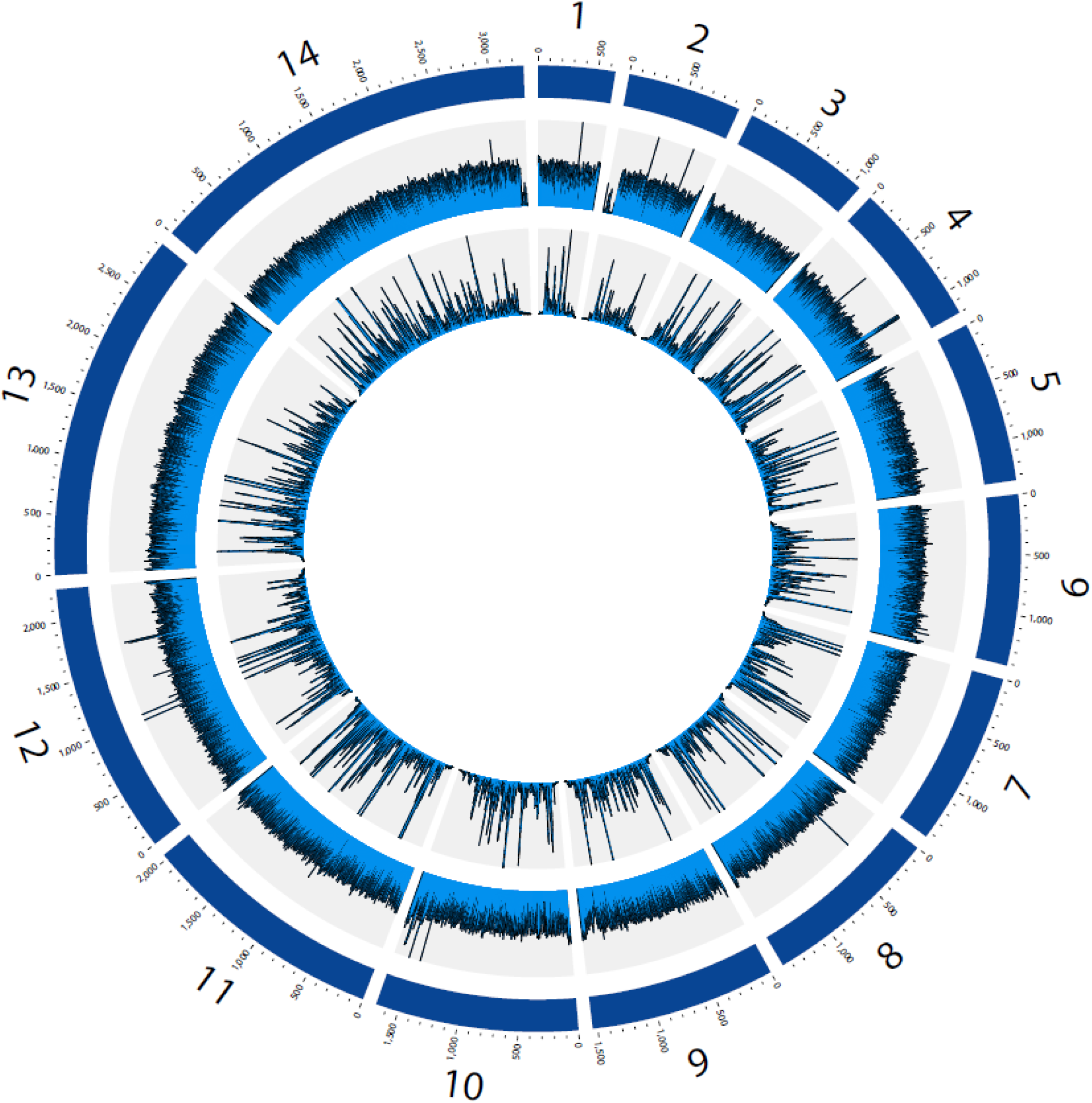
Distribution of the reads along the 14 *P. falciparum* chromosome for M2-1_5F_ and M2_WGA_ mock samples. The three rings represent, from outermost to innermost, the 14 *P. falciparum* chromosomes (illustrated to scale in kb), and the average read depth in 1 kb windows for M2-1_5F_ and for M2_WGA_, respectively. For ease of representation, the maximum depth for M2-1_5F_ and M2_WGA_ was fixed at 200× and 500× respectively.

### Application of 5WBF to *P. falciparum* clinical isolates

We then tested the 5WBF procedure on *P. falciparum* clinical samples with parasitemias ranging from 0.04 to 5.5% (**Table 3**). We used 50 μL (*i.e.* one drop) and 200 μL of whole blood from patients to match blood volumes routinely collected in a clinical context.

**Table 3.**
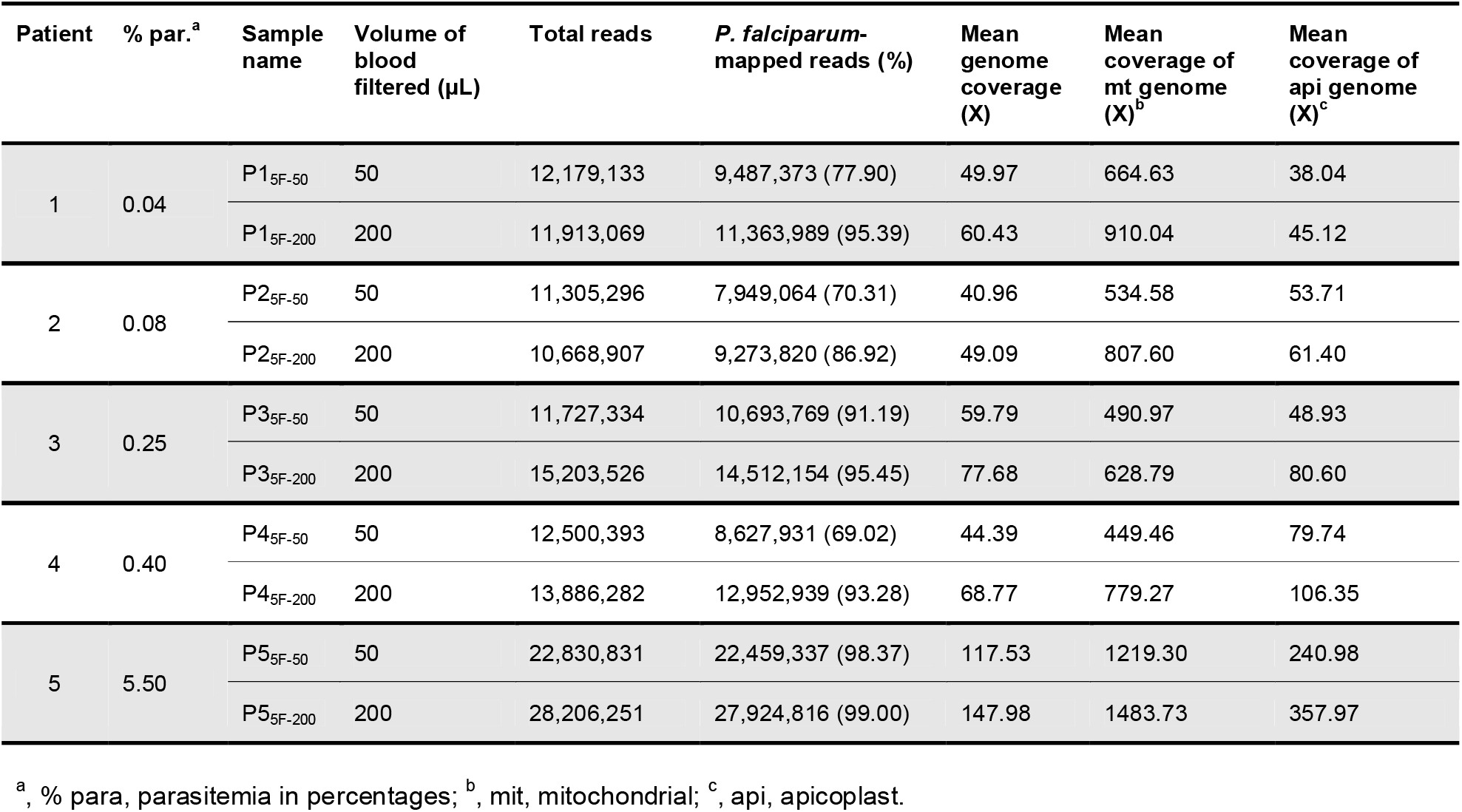
5WBF-based WGS statistics from five *P. falciparum* clinical isolates.

| Patient | % par.<sup>a</sup> | Sample name | Volume of blood filtered (μL) | Total reads | *P. falciparum*-mapped reads (%) | Mean genome coverage (X) | Mean coverage of mt genome (X)<sup>b</sup> | Mean coverage of api genome (X)<sup>c</sup> |
| --- | --- | --- | --- | --- | --- | --- | --- | --- |
| 1 | 0.04 | P1<sub>5F-50</sub> | 50 | 12,179,133 | 9,487,373 (77.90) | 49.97 | 664.63 | 38.04 |
|  |  | P1<sub>5F-200</sub> | 200 | 11,913,069 | 11,363,989 (95.39) | 60.43 | 910.04 | 45.12 |
| 2 | 0.08 | P2<sub>5F-50</sub> | 50 | 11,305,296 | 7,949,064 (70.31) | 40.96 | 534.58 | 53.71 |
|  |  | P2<sub>5F-200</sub> | 200 | 10,668,907 | 9,273,820 (86.92) | 49.09 | 807.60 | 61.40 |
| 3 | 0.25 | P3<sub>5F-50</sub> | 50 | 11,727,334 | 10,693,769 (91.19) | 59.79 | 490.97 | 48.93 |
|  |  | P3<sub>5F-200</sub> | 200 | 15,203,526 | 14,512,154 (95.45) | 77.68 | 628.79 | 80.60 |
| 4 | 0.40 | P4<sub>5F-50</sub> | 50 | 12,500,393 | 8,627,931 (69.02) | 44.39 | 449.46 | 79.74 |
|  |  | P4<sub>5F-200</sub> | 200 | 13,886,282 | 12,952,939 (93.28) | 68.77 | 779.27 | 106.35 |
| 5 | 5.50 | P5<sub>5F-50</sub> | 50 | 22,830,831 | 22,459,337 (98.37) | 117.53 | 1219.30 | 240.98 |
|  |  | P5<sub>5F-200</sub> | 200 | 28,206,251 | 27,924,816 (99.00) | 147.98 | 1483.73 | 357.97 |

<sup>a</sup>, % para, parasitemia in percentages; <sup>b</sup>, mit, mitochondrial; <sup>c</sup>, api, apicoplast.

First, we assessed parasite and human DNA amount by qPCR, expressed in Ct (cycle threshold), before and after 5WBF (**Additional file 1: Table S3**). Slightly higher Ct values were observed after 5WBF for the parasite qPCR assay (Ct_5WBF_ – Ct_unfiltered_: mean = 1.1, min = −1, max = 3; n = 5 samples). In contrast, a dramatic increase in Ct values was observed after 5WBF for the human qPCR assay (Ct_5WBF_ – Ct_unfiltered_: mean = 14, min = 10, max = 16; n = 5 samples). Accordingly, the amount of total DNA quantified by Qubit was drastically lower in the 5WBF-treated samples compared to their unfiltered counterparts (**Additional file 1: Table S3**).

With WGS data, an average of 81.4% (standard deviation: 13.0) and 94.0% (standard deviation: 4.5) of the reads mapped to the *P. falciparum* genome when 50 μL and 200 μL of whole blood were filtered respectively (**Figure 5A** and **Table 3**). The mean genome coverage, including organellar genomes, was systematically higher for 200 μL than 50 μL of whole blood (**Table 3**). However, at the 10× depth threshold the *P. falciparum* genome coverage was similar whether using 50 μL or 200 μL of whole blood (average: 97%; standard deviation: 1.2; **Figure 5B**).

**Figure 5.**
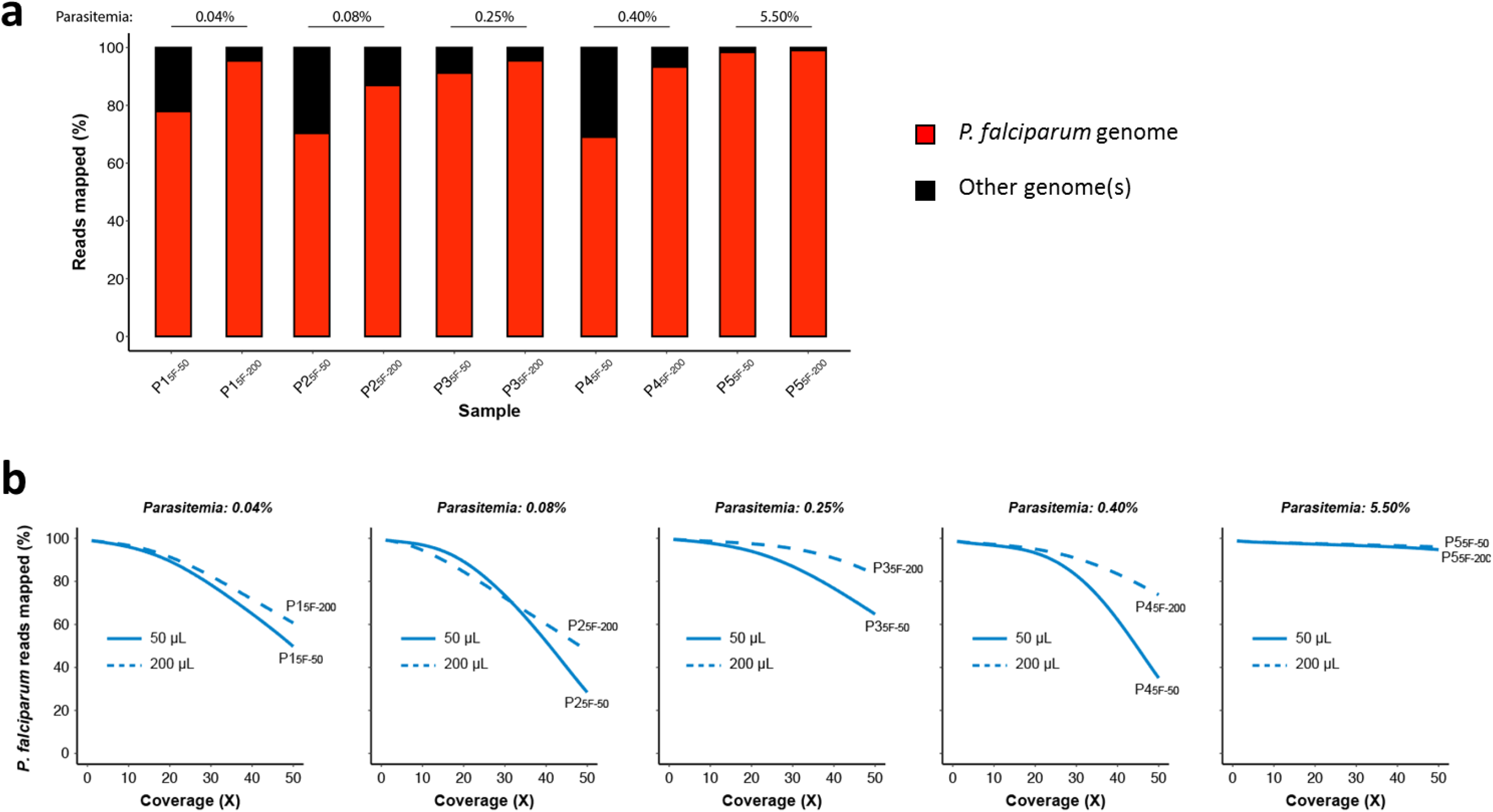
Proportions of mapped reads and *P. falciparum* genome coverage from clinical samples. (**a**) Proportion of reads mapping to the *P. falciparum* nuclear and organellar genomes. Red and black colors respectively indicate the proportion of reads mapping and not mapping to the *P. falciparum* genomes. (**b**) Genome fraction coverage from 1× to 50× depth. Data from 50 μL and 200 μL of filtered whole blood are indicated in solid and dashed lines respectively.

We then explored the two gene-level metrics as previously done for mock samples. We focused on the sample with the lowest parasitemia (0.04%), but with different volume of blood filtered (P1_5F-50_ and P1_5F-200_; **Table 3**). 89.7% (4,899/5,460) and 91.8% (5014/5,460) of *P. falciparum* genes were fully covered at ≥ 10× depth with P1_5F-50_ and P1_5F-200_ respectively, including the major drug resistance genes and all organellar genes (**Figure 6**).

**Figure 6.**
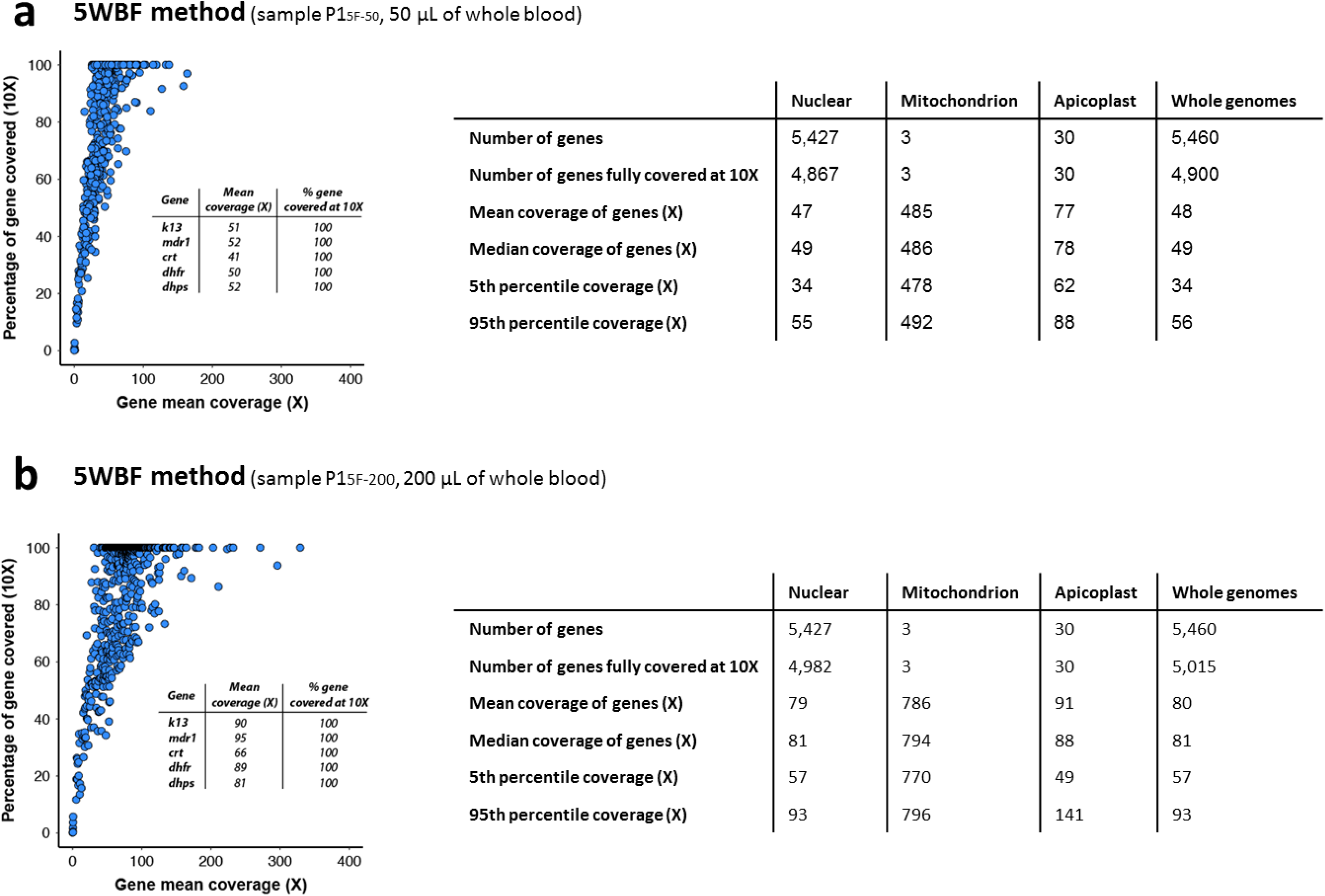
Comparison of gene coverage depth between P1_5F-50_ and P1_5F-200_ 5WBF-treated clinical samples. (**a**) Coverage depth of all genes for P1_5F-50_. Gene distribution based on the percentage of gene covered at ≥ 10× depth. Each blue point corresponds to a gene. Mitochondrial genes were discarded for ease of representation. The *insert* table indicates the mean coverage and the percentage of gene covered at ≥ 10× of five drug resistance genes. Descriptive statistics included the total number of *P. falciparum* (3D7) genes, the number of genes fully covered at ≥ 10×, the mean and median coverage of all genes, and the 5^th^ and 95^th^ percentiles of depth coverage. Genes were partitioned as of either nuclear, mitochondrial, or apicoplast origins. (**b**) Coverage depth of all genes for P1_5F-_200. Description of the plot and the tables are the same as in (a).

Finally, the per-gene copy number was compared for P1_5F-50_ and P1_5F-200_. Beforehand, all the variant surface antigens gene families (*var*, *stevor*, *rifin*, *phist* and *Plasmodium exported protein-*encoding genes) were removed to avoid any bias in the analysis (4,816 remaining genes). Similar profiles were obtained for P1_5F-50_ and P1_5F-200_ and no gene amplification was detected (Spearman’s rank correlation: *p* < 0.001, r□=□0.72; **Figure 7**). Among the other samples tested, P5_5F-50_ and P5_5F-200_ harbored three copies of the *GTP cyclohydrolase 1* gene (*gch1*; PF3D7_1224000) and of the four neighboring genes (PF3D7_1223700, PF3D7_1223800, PF3D7_1223900 and PF3D7_1224100; **Additional file 1: Figure S1**). The total amplicon size was 10.5 kb and resembled to the one detected in Thai isolates [19]). Altogether, 50 μL of whole blood at 0.04% parasitemia treated by 5WBF permitted to explore per-gene copy number in a clinical context.

**Figure 7.**
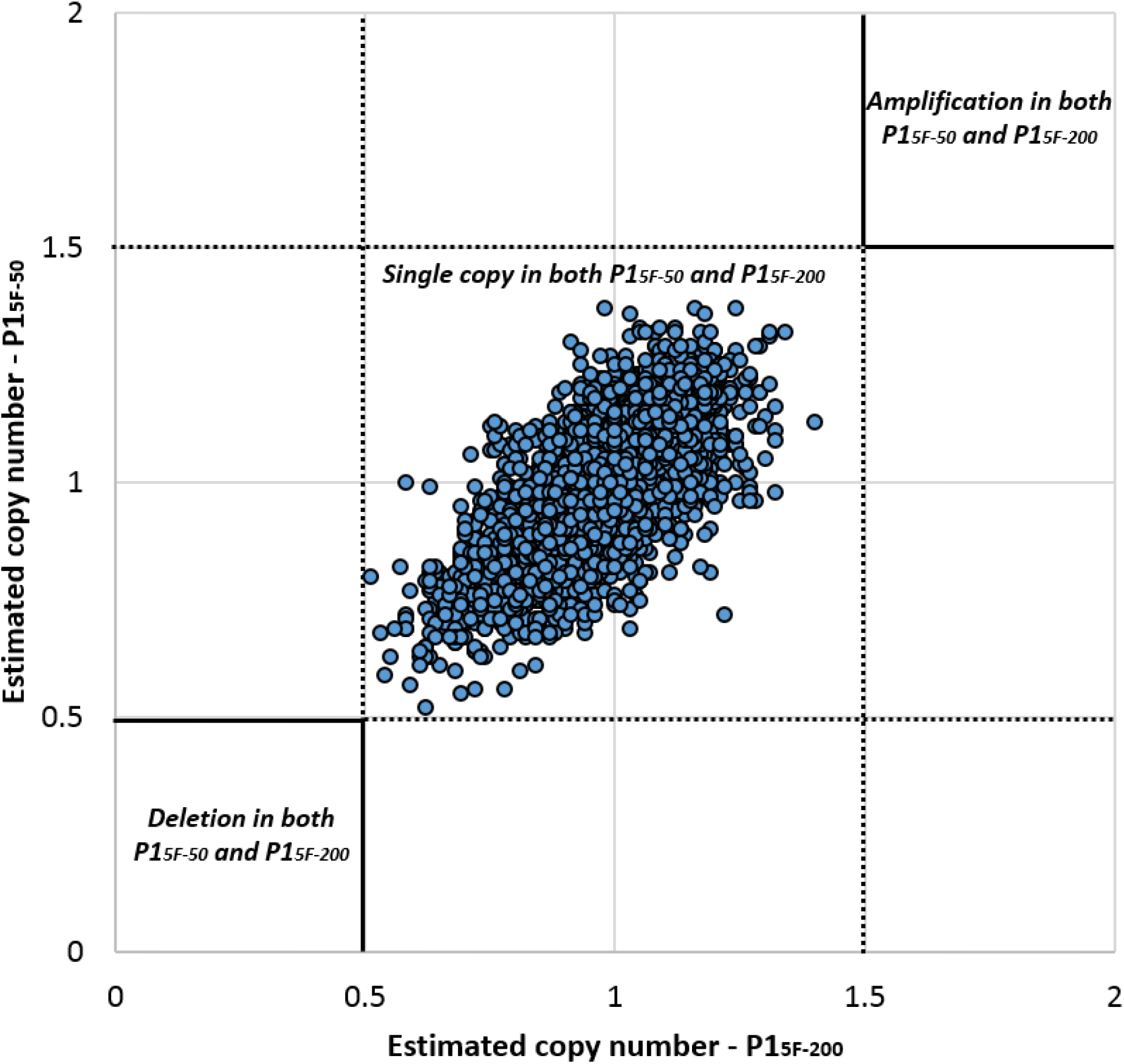
Estimation of per-gene copy number for P1_5F-50_ and P1_5F-200_ clinical samples using PlasmoCNVScan. Each point corresponds to a gene. A value < 0.5 suggests a gene deletion, while a value > 1.5 suggests a gene amplification. Values between 0.5 and 1.5 are supposed to be single gene copy. A positive correlation was observed between the two samples (Spearman’s rank correlation: p < 0.001, r□=□0.72).

## DISCUSSION

The ability to produce high-quality sequencing data from *P. falciparum* clinical samples has valuable implications for public health. The sWGA method has massively facilitated the generation of WGS from clinical whole blood samples stored on DBS. However, sWGA-based methods present several drawbacks. Current primers used to selectively amplify the *P. falciparum* genome leads to the nearly complete loss of the mitochondrial and apicoplast genomes [9]. Furthermore, a large proportion of reads often do not map to the *P. falciparum* genome, suggesting that some contaminant human DNA remain after the sWGA step [12]. Finally, reads mapping to the reference genome are not homogeneously distributed across the genome, precluding any investigation based on read distribution such as the measure of per-gene copy number [18]. Here, we demonstrate the usefulness of 5WBF, a new leucodepletion protocol based on commercial 5 μm filters. Other filtration approaches were already successfully developed but also have their own drawbacks – whether in terms of costs, blood volumes, or standardization [7,8]. Here, we extensively used 5WBF for WGS purposes, but this strategy may likely be useful for other sequencing application such as RNAseq, which often suffers from contaminant human DNA when applied to *P. falciparum* isolates and thus requiring additional costs.

Sequencing data obtained with whole blood samples treated by 5WBF revealed that all the three *P. falciparum* genomes (nuclear, mitochondrial and apicoplast) were covered with high depth coverage. Almost all of the *P. falciparum* genes were fully covered at ≥ 10× depth, except the highly variable *var* and *rifin* gene families. Capturing the organellar genomes is especially important since they carry drug resistance genes such as *cytb* or *rps4* [20–22] or can inform on the geographic origin and evolution of the parasites [23,24]. Finally, the homogeneous distribution of reads across the genome makes it possible to detect robustly copy number variations. Such a result is very relevant since amplification of *mdr1* and *plasmepsin 2/3* modulates the resistance to multiple antimalarials [25–30].

The 5WBF procedure was extensively tested here using the 5 μm Minisart NML® syringe filter from the manufacturer Sartorius. Other commercially available 5 μm might also be suitable and would need validation experiments. Financially, the cost of the 5 μm Minisart NML® syringe filter varies from 1.0 to 1.7€ per unit depending on suppliers. This is about 10 times cheaper than the Plasmodipur filter (Europroxima, Arnhem, The Netherlands, Cat. 8011Filter25U [6,8]). Similarly, sWGA-based methods are more expensive than 5WBF since Phi29 DNA polymerase, primer sets and subsequent purification with Agencourt Ampure XP beads increase the cost to approximately 6-8€ per sample [9–11]. Only the leucodepletion-based method through CF11 cellulose column has a similar implementation cost [7]. However, these are homemade columns, and thus requires an extended preparation time and likely lack the standardized and certified quality of commercially available filters. Altogether, the 5WBF procedure provides remarkable add-ons: simplicity and speed of the filtration procedure, standardized and ready-to-use filters, low cost per sample, and high quality of WGS data.

The principal limitation of the 5WBF procedure is that, as for any filtration procedure, it introduces practical constraints related to centrifugation of the resulting filtrate to pellet RBCs (or decantation, if no power available as in [8]). Also, 5WBF-treated blood samples could be stored after centrifugation as DBS on filter papers, as previously done with Plasmodipur filters [8], but this needs to be tested.

The 5WBF+WGS procedure described here was successfully tested and replicated with several *P. falciparum* blood samples exhibiting various parasitemias at the ring stage. Whether this technique works at parasitemias lower than 0.02%, with RBCs infected with more mature *P. falciparum* stages (trophozoite and schizont), and with other *Plasmodium* species needs to be rigorously tested. For example, our initial attempts on *P. ovale* clinical samples and on *in vitro P. falciparum* culture containing more mature asexual stages (trophozoite and schizont) indicated highly variable results regarding the capacity of these other species and stages to pass through the 5 μm filter we used (data not shown). Considering that both the size and deformability of the host RBC vary according to the infecting *Plasmodium* species [31,32] and stage maturity, a 5 μm filter might not be suitable for other *Plasmodium* species infecting humans.

## CONCLUSION

In summary, we report a simple and cheap filtration procedure – 5WBF – that efficiently depletes human leukocytes from minute amounts of human ring-stage *P. falciparum*-infected blood. The highly enriched *P. falciparum* genomic DNA obtained is suitable for high-quality, extensive WGS and downstream analyses, including coverage of organellar genomes and detection of copy number variations.

## LIST OF ABBREVIATIONS

5WBF: 5 μm Whole Blood Filtration
Ct: Cycle threshold
DBS: Dried blood spots
RBCs: Red blood cells
sWGA: Selective whole-genome amplification
WGS: Whole-genome sequencing

## DECLARATION

### Ethics approval

No institutional review board approval was required according to French legislation (article L. 1111-7 du Code de la Santé Publique, article L. 1211-2 du Code de Santé Publique, articles 39 et suivants de la loi 78-17 du 6 janvier 1978 modifiée en 2004 relative à l’informatique, aux fichiers, et aux libertés). Samples received at the French Malaria Reference Center, (Paris, France) were registered and declared for research purposes as a biobank for both the Assistance Publique des Hôpitaux de Paris and Santé Publique France.

### Consent for publication

There are no case presentations that require disclosure of respondent’s confidential data/information in this study.

### Availability of data and materials

The script used to calculate the percentage of each *P. falciparum* gene covered at ≥ 10× depth and the per-gene mean coverage depth was deposited on github: https://github.com/Rcoppee/Scan_gene_coverage. The datasets analyzed in the study are available from the corresponding authors on request.

### Author’s contributions

RC, FA, SH, and JC conceived and coordinated the study. AM, VS, CK, and LP performed the filtrations. RC performed sWGA. SH and VS participated in sample collection. LA and FL performed next-generation sequencing. RC performed data analyses and script production. RC drafted the manuscript. RC, AM, VS, CK, FA, SH and JC participated in the editing and final preparation of the manuscript. All authors read and approved the final manuscript.

