## Additional File 1 for "5WBF: A low-cost and straightforward whole blood filtration method suitable for whole-genome sequencing of *Plasmodium falciparum* clinical isolates"

**Additional file 1 : Supplementary Figures and Tables**

This supplemental file has been provided by the authors to give readers additional information about their work.

**Figure S1 ……………………….…………………………………………………………………….. 2**

**Table S1 …………………………………………………………………………………….…..…….. 3**

**Table S2 …………………………………………………………………………………….…..…….. 3**

**Table S3 …………………………………………………………………………………….…..…….. 3**

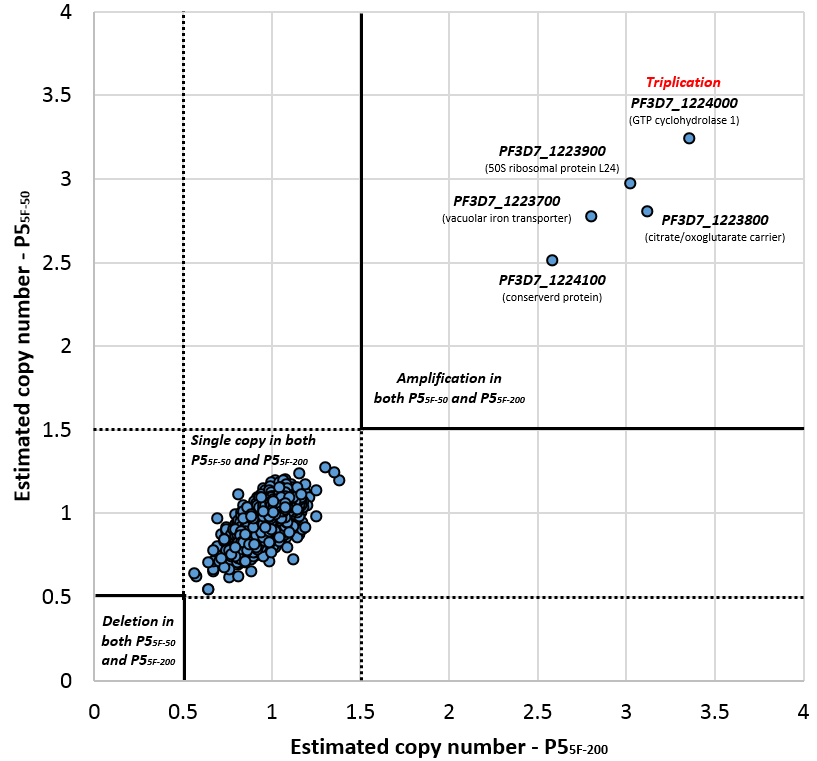

**Figure S1 – Estimation of per-gene copy number for P5_5F-50_ and P5_5F-200_ using PlasmoCNVScan**. Each point corresponds to a gene. A value < 0.5 suggests a gene deletion, while a value > 1.5 suggests a gene amplification. Values between 0.5 and 1.5 are supposed to be single gene copy. *gch1* (PF3D7_1224000) and four neighboring genes (PF3D7_1223700, PF3D7_1223800, PF3D7_1223900 and PF3D7_1224100) have values > 2.5, suggesting a triplication.

**Table S1 – Clinical information of the patients included in the study.**

| **Patient** | **Sex** | **Age** | **Infection country** | **Prophylaxis** | **% par. ^a^** | **Volume of blood filtered (µL)** | **Sample** |
| --- | --- | --- | --- | --- | --- | --- | --- |
| 1 | M | 20 | Ivory Coast | None | 0.04 | 50 | P1_5F-50_ |
|  |  |  |  |  |  | 200 | P1_5F-200_ |
| 2 | M | 44 | Ivory Coast | NA | 0.08 | 50 | P2_5F-50_ |
|  |  |  |  |  |  | 200 | P2_5F-200_ |
| 3 | M | 45 | Cameroon | NA | 0.25 | 50 | P3_5F-50_ |
|  |  |  |  |  |  | 200 | P3_5F-200_ |
| 4 | M | 44 | Central African Republic | None | 0.4 | 50 | P4_5F-50_ |
|  |  |  |  |  |  | 200 | P4_5F-200_ |
| 5 | M | 29 | Ivory Coast | None | 5.5 | 50 | P5_5F-50_ |
|  |  |  |  |  |  | 200 | P5_5F-200_ |

Note – NA, Not Available. ^a^ % par., parasitemia in percentages.

**Table S2 – sWGA primers for *P. falciparum.***

| **Primer name** | **Primer sequence** |
| --- | --- |
| Pf1 | ATATATATAT*A |
| Pf2 | TATATATATAT*T |
| Pf3 | TATATATATA*A |
| Pf4 | TAATATATA*T |
| Pf5 | TATATATATT*T |
| Pf6 | ATTATTATTA*T |
| Pf7 | TAATAATAAT*A |
| Pf8 | AAAAAAAAAAA*A |
| Pf9 | AATAATAATA*A |
| Pf10 | TATTATATA*T |

* phosphorothioate bond.

**Table S3 – Content in *P. falciparum* species and total DNA before and after 5WBF measured by Qubit and qPCR DNA quantification.**

| **Patient** | **% par. ^a^** | **Sample** | **Before/after**  **5WBF** | **Qubit total DNA ^b^** | **Vol. of blood filtered (µL)** | ***H. sapiens* ΔCt ^c^** | ***P. falciparum* ΔCt ^d^** |
| --- | --- | --- | --- | --- | --- | --- | --- |
| 1 | 0.04 | P1_5F-50_ | Before | 2.07 | 50 | 16 | 1 |
|  |  |  | After | < 0.01 |  |  |  |
|  |  | P1_5F-200_ | Before | 12.1 | 200 | 14 | 1 |
|  |  |  | After | 0.05 |  |  |  |
| 2 | 0.08 | P2_5F-50_ | Before | 2.95 | 50 | 12 | 2 |
|  |  |  | After | 0.02 |  |  |  |
|  |  | P2_5F-200_ | Before | 20.4 | 200 | 14 | 1 |
|  |  |  | After | 0.09 |  |  |  |
| 3 | 0.25 | P3_5F-50_ | Before | 7.80 | 50 | 16 | 3 |
|  |  |  | After | 0.02 |  |  |  |
|  |  | P3_5F-200_ | Before | 19.30 | 200 | 15 | 1 |
|  |  |  | After | 0.30 |  |  |  |
| 4 | 0.40 | P4_5F-50_ | Before | 0.10 | 50 | >10 | 2 |
|  |  |  | After | < 0.01 |  |  |  |
|  |  | P4_5F-200_ | Before | 9.52 | 200 | 15 | 2 |
|  |  |  | After | 0.10 |  |  |  |
| 5 | 5.50 | P5_5F-50_ | Before | 2.39 | 50 | 15 | -1 |
|  |  |  | After | 0.22 |  |  |  |
|  |  | P5_5F-200_ | Before | 13.40 | 200 | 13 | -1 |
|  |  |  | After | 2.23 |  |  |  |

Note – ^a^ % par., parasitemia in percentages. ^b^ Total DNA was quantified using Qubit® dsDNA high sensitivity (Thermo Fisher Scientific). ^c^ Ct, cycle threshold; ΔCt = Ct_5WBF_ – Ct_unfiltered_. ^d^ Ct, cycle threshold; ΔCt = Ct_5WBF_ – Ct_unfiltered_.
